# Maternal Obesity-Induced Endoplasmic Reticulum Stress Causes Metabolic Alterations and Abnormal Hypothalamic Development in the Offspring

**DOI:** 10.1101/637066

**Authors:** Soyoung Park, Alice Jang, Sebastien G. Bouret

## Abstract

The steady increase in the prevalence obesity and associated type II diabetes is a major health concern, particularly among children. Maternal obesity represents a risk factor that contributes to metabolic perturbations in the offspring. Endoplasmic reticulum (ER) stress has emerged as a critical mechanism involved in leptin resistance and type 2 diabetes in adult individuals. Here, we used a mouse model of maternal obesity to investigate the importance of early life ER stress in the nutritional programming of metabolic disease. Offspring of obese dams displayed increased body weight, adiposity, food intake and developed glucose intolerance. Moreover, maternal obesity disrupted the development of melanocortin circuits associated with neonatal hyperleptinemia and leptin resistance. ER stress-related genes were upregulated in the hypothalamus of neonates born to obese mothers and neonatal treatment with the ER stress-relieving drug tauroursodeoxycholic acid improved metabolic and neurodevelopmental deficits and reverses leptin resistance in neonates born to obese dams.

## Introduction

A major shift in our nutritional environment has greatly contributed to the recent obesity epidemic. There is growing evidence that adverse fetal and early postnatal environments increase the risk of developing obesity. In particular, accumulative evidence from both human and animal studies demonstrated that exposure to maternal obesity predisposes offspring to develop obesity and other metabolic dysfunctions later in life [1,2,3]. The hypothalamus is involved in the control of food intake and energy expenditure and is a prime target of developmental the programming of obesity induced by maternal and perinatal nutritional imbalances [1,2,4,5,6,7]. A primary importance has been given to the arcuate nucleus of the hypothalamus (ARH) because it contains two main neuronal populations that play a major role in energy homeostasis: the anorexigenic pro-opiomelanocortin (POMC)-expressing neurons and the orexigenic neuropeptide Y (NPY) and agouti-related peptide (AgRP)-expressing neurons. The adipocyte-derived hormone leptin directly targets these neuronal populations to cause weight loss effects by stimulating POMC neurons and inhibiting AgRP/NPY neurons. Leptin also promotes the development of POMC and AgRP/NPY axonal projections during early postnatal life [8]. Prior studies have shown that maternal obesity disrupts the normal development of these neuronal circuits [9,10,11]. However, the cellular mechanisms involved in hypothalamic development and how these mechanisms are perturbed in a context of maternal obesity remain elusive.

Endoplasmic reticulum (ER) stress provides an attractive mechanism to underlie the programming effects of maternal obesity. Alterations in cellular homeostasis can lead to ER stress and the activation of the unfolded protein response (UPR) pathway. Previous studies have demonstrated that ER stress and UPR signaling pathway activation play important roles in obesity-induced insulin resistance and type 2 diabetes during adult life. Obesity caused by leptin deficiency or high-fat feeding in mice induces ER stress in peripheral tissues as well as in the hypothalamus [12,13,14]. Furthermore, relieving ER stress with chemical chaperones, *i.e.*, agents that have the ability to increase ER folding machinery, increases insulin sensitivity and reverses type 2 diabetes in adult *ob/ob* mice and improves leptin sensitivity in adult obese mice fed a high-fat diet (HFD) [13,14]. Moreover, genetic manipulation of the unfolded protein response transcription factor spliced X-box binding protein (Xbp1) specifically in POMC neurons protects against diet-induced obesity and ameliorates leptin and insulin sensitivity [15]. Despite accumulative evidence supporting a role for ER stress in metabolic regulation, an association between maternal obesity, ER stress, and the programming of obesity and hypothalamic development has not yet been established.

In the present study, we investigated whether maternal diet-induced obesity induces ER stress during neonatal life in the offspring and how it contributes to the nutritional programming of obesity and hypothalamic development. We found that maternal obesity causes metabolic and neurodevelopmental alterations in the offspring accompanied with elevated ER stress in the hypothalamus and pancreas during postnatal development. Moreover, we report that pharmacological inhibition of ER stress has long-term beneficial effects on body weight, body composition, energy balance, glucose homeostasis, leptin sensitivity, and POMC axonal projections in the offspring born to obese dams. Finally, our study reveals that the neurodevelopmental effects of maternal obesity likely involve direct inhibitory action of saturated fatty acids on arcuate axon growth.

## Results

### Maternal obesity causes metabolic disturbances in the offspring

A mouse model of maternal obesity induced by high-fat high-sucrose (HFHS) feeding during pregnancy and lactation was used to study the effects of maternal obesity on the offspring’s metabolism and development. Adult female mice were either fed a HFHS (58% kcal fat w/ sucrose) or a control diet (6% calories from fat) six weeks before breeding. Dams were kept on their respective diet throughout pregnancy and lactation. A significant increase in dams’ weight gain was observed as early as 4 weeks after HFHS diet began and persisted throughout the HFHS exposure (Fig 1A). This elevated body weight was associated with increased fat mass (Fig 1B). Moreover, dams fed a HFHS diet displayed altered glucose tolerance during gestation (Fig 1C).

**Fig 1.**
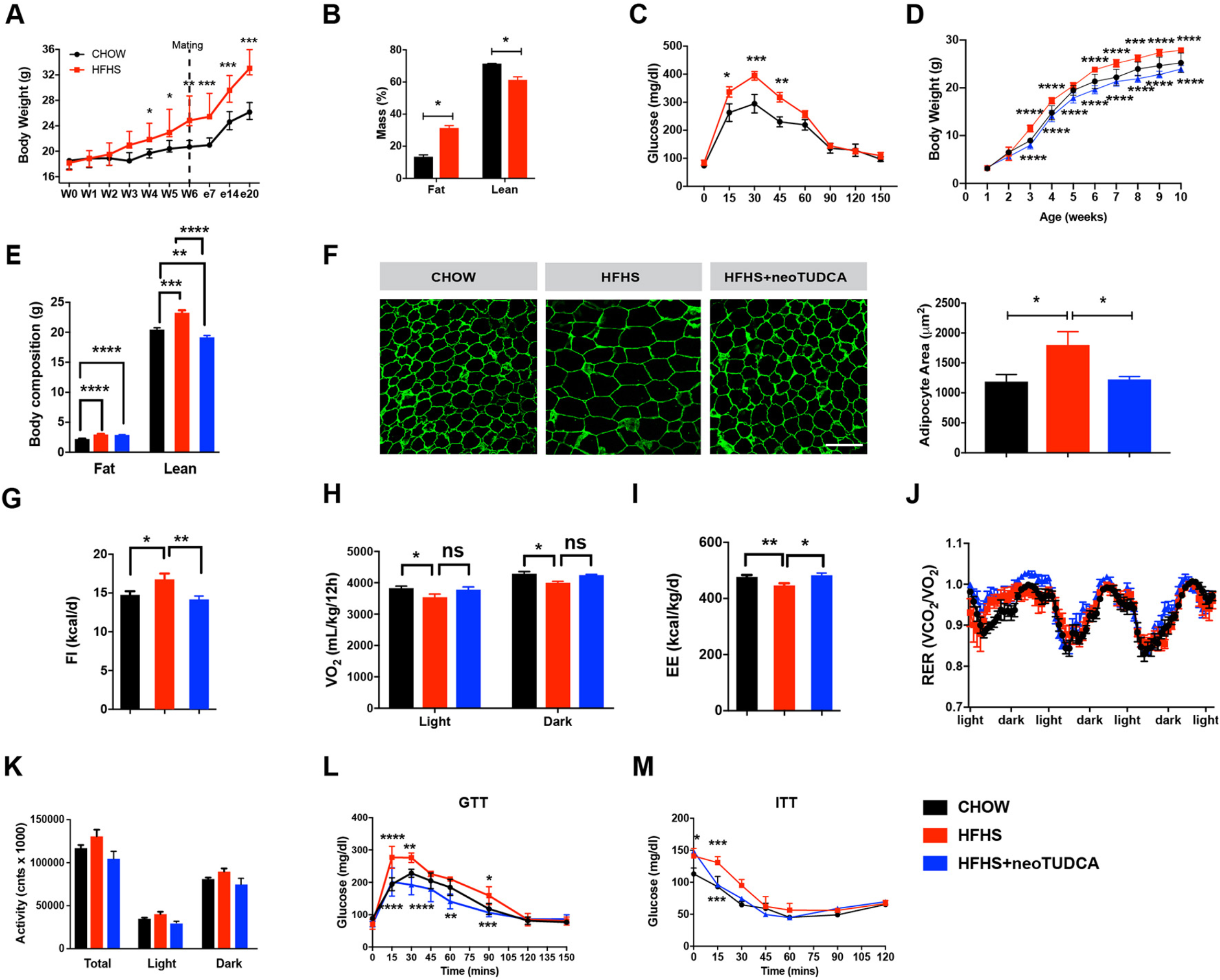
Maternal obesity impairs energy balance and glucose homeostasis in the offspring and neonatal TUDCA treatment improves this metabolic malprogramming. (A) Body weight curves of adult female mice fed a chow or a high-fat high-sucrose (HFHS) diet before and during pregnancy (n = 5 per group). (B) Body composition (n = 3 per group) and (C) glucose tolerance test (GTT) (n = 4-5 per group) of pregnant female mice fed a chow or HFHS diet at gestational day 16. (D) Body weight curves of mice born to chow-fed dams, HFHS-fed dams, or HFHS-fed dams and treated with tauroursodeoxycholic acid (TUDCA) neonatally (neoTUDCA) (n = 5-10 per group). (E) Average body composition (n = 5-8 per group) and (F) representative images and quantification of adipocyte size (immunostained for perilipin, *green* fluorescence) of 10-week-old mice born to chow-fed dams, HFHS-fed dams, or HFHS-fed dams and treated with TUDCA neonatally (n =4-5 per group). (G) Food intake, (H) oxygen consumption, (I) energy expenditure, (J) respiratory exchange ratio (RER), and (K) locomotor activity of 10-week-old mice born to chow-fed dams, HFHS-fed dams, or HFHS-fed dams and treated with TUDCA neonatally (n = 3-8 per group). (L) Glucose (GTT) and (M) insulin tolerance tests (ITT) of 7-8-week-old mice born to chow-fed dams, HFHS-fed dams, or HFHS-fed dams and treated with TUDCA neonatally (n = 4-8 per group). Data are presented as mean ± SEM. \**P* ≤ 0.05, \*\**P* ≤ 0.01, \*\*\**P* ≤ 0.001, and \*\*\*\**P* ≤ 0.0001 *versus* chow groups. Statistical significance between groups was determined by unpaired two-tailed Student’s *t* test (E), one-way ANOVA (F, I, K), and two-way ANOVA (A-D, H, L, M) followed by Tukey’s Multiple Comparison test. Scale bar, 100 µm.

The offspring of HFHS-fed dams had heavier body weights at weaning and this elevated body weight persisted into adulthood (Fig 1D). We also evaluated body composition and found that adult animals born to obese dams displayed elevated fat and lean mass compared to control mice (Fig 1E). Moreover, neonatal exposure to HFHS caused adipocyte hypertrophy as revealed by a 1.5-fold increase in adipocyte size in epididymal white adipose tissue (Fig 1F). There was also an increase in food intake, and decreases in oxygen consumption (VO_2_) and energy expenditure in adult animals born to obese dams (Fig 1G-I). Respiratory exchange ratio and locomotor activity were not significantly different compared to controls (Fig 1J and K). However, adult mice born to obese dams displayed impaired glucose and insulin tolerances compared to mice born to lean dams (Fig 1L and M).

### Maternal obesity induces ER stress during postnatal development and neonatal TUDCA treatment has long-term beneficial metabolic effects

To examine if maternal obesity was associated with activation of ER stress response in the offspring, we measured the expression levels of the following ER stress markers in metabolically-relevant tissues: activating transcription factor 4 (*Atf4)*, 6 (*Atf6)*, X-box binding protein (*Xbp1)*, glucose regulated protein GRP78 (referred to as *Bip*), and CCAAT-enhancer-binding protein homologous protein (*Chop*), in P10 and adult animals born to chow- or HFHS-fed dams. The mRNA levels of *Atf4*, *Atf6, Xbp1*, *Bip*, and *Chop* were significantly elevated in the ARH of P10 mice born to obese dams (Fig 2A). Moreover, expression of *Atf4*, *Atf6, Xbp1*, and *Chop* mRNAs were significantly higher in the ARH of adult mice born to HFHS-fed dams (Fig 2B). In contrast, only *Atf4* mRNA was significantly increased in the paraventricular nucleus (PVH) of P10 mice (Fig 2C and D). *Atf4, Atf6*, and *Xbp1* mRNAs were elevated in the pancreas of P10 pups of HFHS-fed dams (Fig 2E), but these markers were not significant changed in neonatal liver and fat tissues (Fig 2G and I). In addition, *Atf4, Atf6, Xbp1, and Chop* mRNA levels were higher in the pancreas of adult mice born to obese dams (Fig 2F), but only *Xbp1* and *Xbp1* as well as *Chop* mRNAs were significantly elevated in the liver and fat tissues, respectively, of adult mice of HFHS-fed dams (Fig 2 H and J).

**Fig 2.**
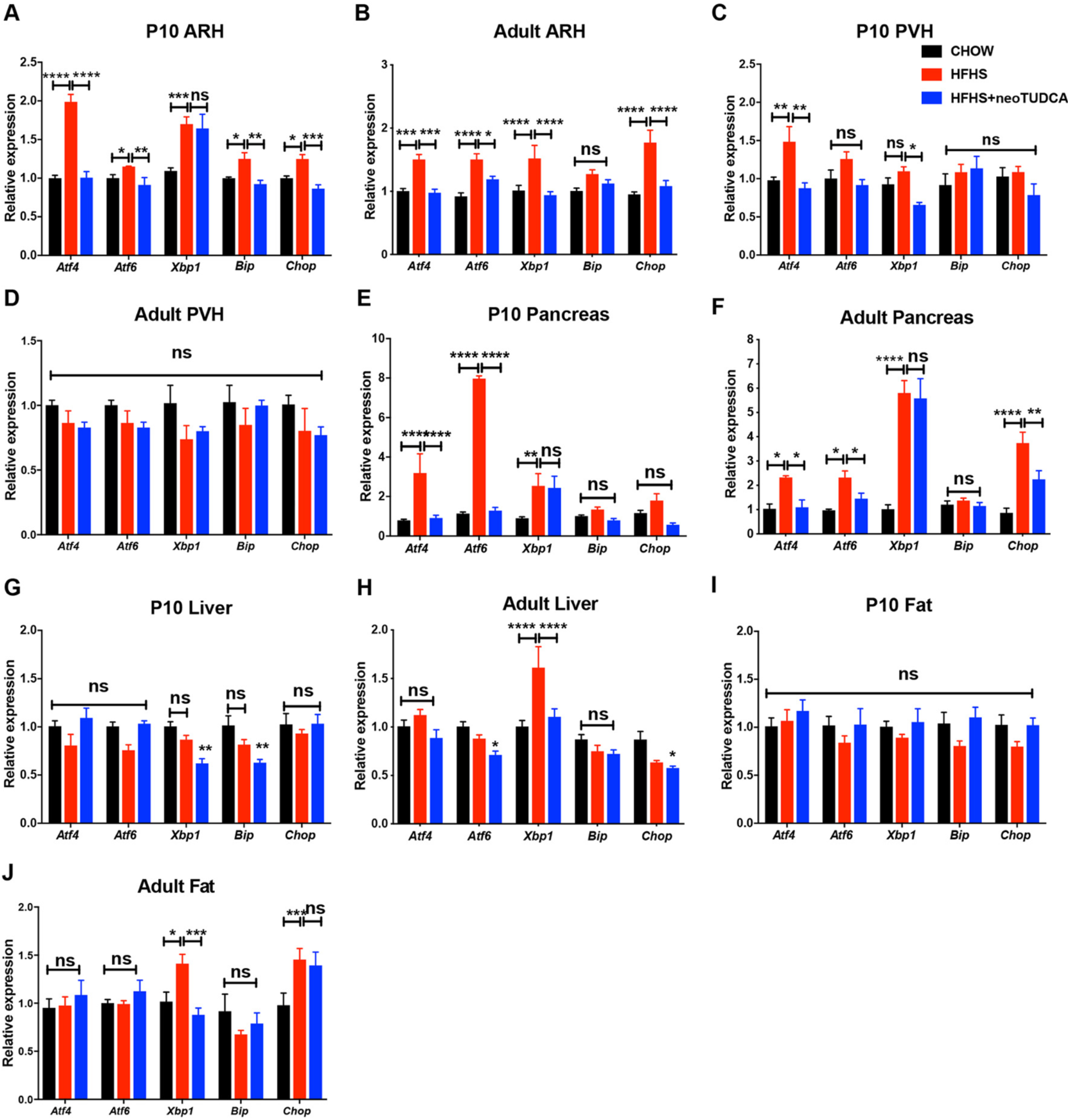
Neonatal TUDCA treatment reverses the elevated expression of ER stress markers in the offspring of obese dams. **(A-J)** Relative expression of *Atf4, Atf6, Xbp1, Bip*, and *Chop* mRNA in (A, B) the arcuate nucleus (ARH), (C, D) paraventricular nucleus (PVH), (E, F) pancreas, (G, H) liver, and (I, J) fat depot of (A, C, E, G, and I) P10 and (B, D, F, H, and J) 10-week-old adult mice born to chow-fed dams, high-fat high-sucrose (HFHS)-fed dams, or HFHS-fed dams and treated with TUDCA neonatally (n = 4-6 per group). Data are presented as mean ± SEM. \**P* ≤ 0.05, \*\**P* < 0.01, \*\*\**P* ≤ 0.001, and \*\*\*\**P* ≤ 0.0001 *versus* other groups. Statistical significance between groups was determined by two-way ANOVA (A-J) followed by Tukey’s Multiple Comparison test.

To investigate the importance of early life ER stress, we treated pups born to HFHS-fed dams with daily peripheral injections of tauroursodeoxycholic acid (TUDCA) from P4 to P16, which represents a critical period for growth and development, including of the hypothalamus [2]. TUDCA is a chemical chaperone of low molecular weight that is well-known to alleviate ER stress [14,16]. Neonatal treatment with TUDCA in animals born to obese dams reversed induction of most ER stress markers in the postnatal and adult ARH, pancreas, liver, and adipose tissue (Fig 2A-J), with the exception of *Xbp1* in P10 ARH and pancreas (Fig 2A and E) and in adult pancreas (Fig 2F), and *Chop* in adult adipose tissue (Fig 2J). Neonatal TUDCA treatment also reduced normal mRNA levels of *Xbp1* in P10 PVH and liver (Fig 2C and G), *Bip* in P10 liver (Fig 2G), and *Atf6* in adult liver (Fig 2H). Physiologically, neonatal TUDCA treatment in animals born to obese dams reversed alterations in body weights, body composition, adipocytes, food intake, energy expenditure, and glucose and insulin tolerances (Fig 1D-G, I, and L-M), with only VO_2_ not being improved (Fig 1H).

Together, these data indicate that maternal obesity causes elevated ER stress levels in metabolically-relevant tissues during postnatal and adult life and that this induction of ER stress is reversible upon neonatal TUDCA treatment, which also causes long-term beneficial effects on energy metabolism.

### Maternal obesity causes hyperleptinemia and reduces hypothalamic leptin sensitivity in the offspring that is reversible with neonatal TUDCA treatment

Previous studies have reported that during perinatal life leptin exerts marked neurodevelopmental and metabolic effects [8,17,18,19,20]. We therefore measured circulating leptin levels in animals exposed to maternal obesity. Maternal HFHS feeding was associated with a marked increase in serum leptin levels in dams at gestational day 16 and in E16.5 embryos (Fig 3A). Serum leptin levels were also elevated in P10 pups born to obese dams, which were normalized upon neonatal TUDCA treatment (Fig 3A). However, serum leptin levels were unchanged in adult mice born to HFHS-fed mothers (Fig 3A). Because leptin’s neurotrophic effects require intact ARH LepRb->pSTAT3 signaling [21], we also evaluated the number of pSTAT3-immunoreactive neurons after peripheral leptin injection and found that leptin treatment resulted in significantly fewer pSTAT3-positive cells in the ARH of P14 pups from obese dams and that neonatal TUDCA treatment enhanced ARH leptin-induced pSTAT3 (Fig 3B). To determine whether maternal obesity affected leptin sensitivity in other hypothalamic nuclei, we also examined leptin-induced pSTAT3-immunoreactivity in the DMH and found that the number of pSTAT3-positive cells was unaltered in the DMH of pups born to HFHS-fed dams (Fig 3B).

**Fig 3.**
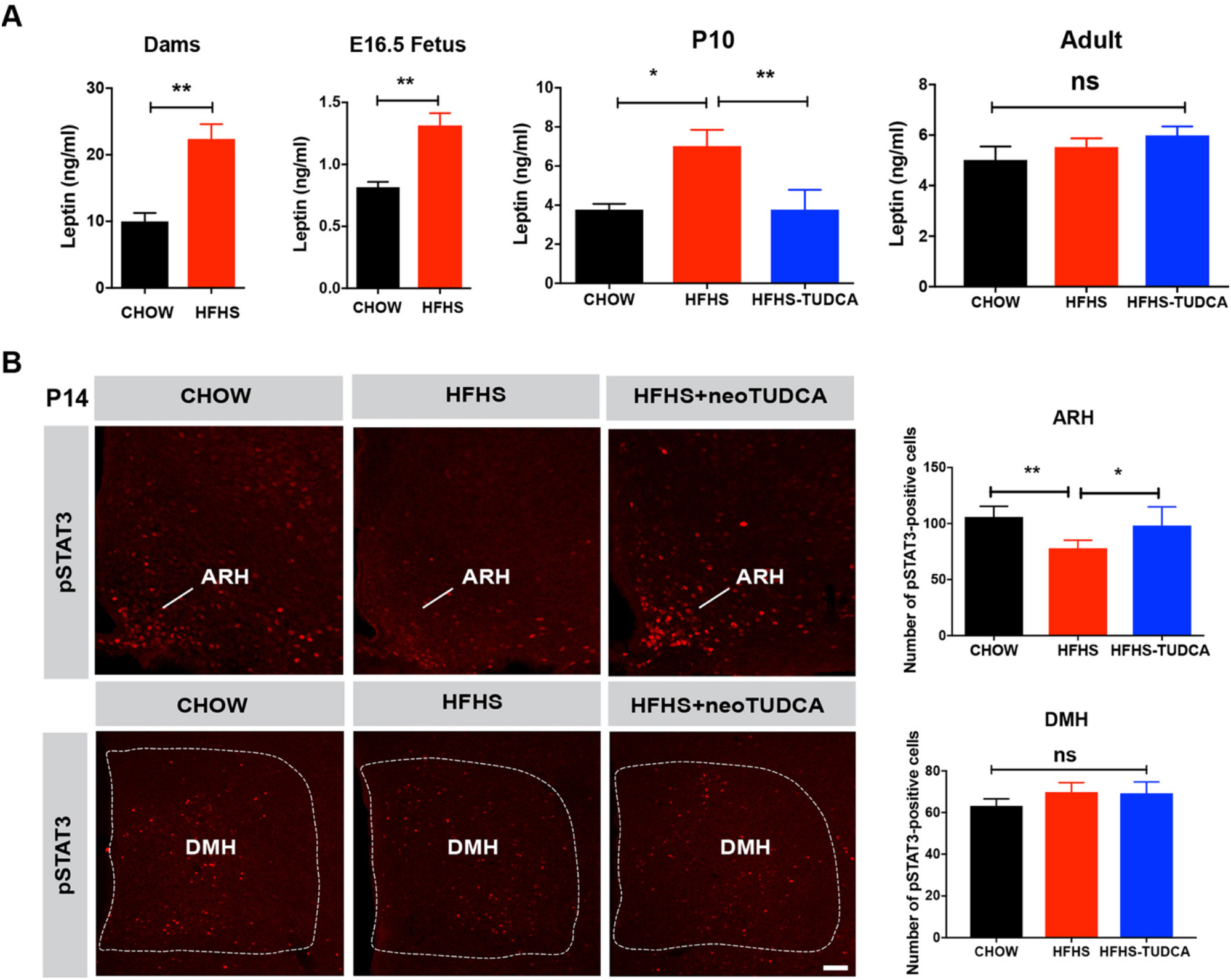
Maternal obesity causes neonatal hyperleptinemia and attenuated response to leptin that can be reversed by neonatal TUDCA treatment. (A) Serum leptin levels in dams at gestational day 16 and E16.5 fetuses of dams fed a chow or HFHS diet and in P10 and 10-week-old mice born to chow-fed dams, high-fat high-sucrose (HFHS)-fed dams, or HFHS-fed dams and treated with TUDCA neonatally (n = 4-8 per group). (B) Confocal images and quantification of the number of leptin-induced pSTAT3-immunoreactive cells in the arcuate nucleus (ARH) and dorsomedial nucleus (DMH) of P14 pups born to chow-fed dams, HFHS-fed dams, or HFHS-fed dams and treated with TUDCA neonatally (n = 5 per group). Data are presented as mean ± SEM. \**P* ≤ 0.05 and \*\**P* < 0.01 *versus* chow groups. Statistical significance was determined by unpaired two-tailed Student’s t test (A), and one-way ANOVA followed by Tukey’s Multiple Comparison test (B). Scale bar, 100 µm.

### Neonatal TUDCA exposure restores disrupted POMC axonal projections in the offspring of HFHS-fed dams

During postnatal development, neuronal projections from the ARH reach their target nuclei, including the PVH, under the influence of leptin and leptin receptor signaling [8,21]. Because our results indicate that maternal obesity alters offspring’s leptin levels and ARH leptin signaling, we next investigated whether maternal obesity disrupts the development of ARH circuits by examining POMC and AgRP neuronal projections, two arcuate neuropeptidergic systems playing a critical role in energy balance. The density of POMC-immunoreactive fibers in the PVH of P14 mice born to obese dams was 2-fold lower than that observed in control mice (Fig 4A). In contrast, the density of AgRP-labeled projections innervating the PVH appeared normal in P14 pups born to HFHS-fed dams (Fig 4A). Also, the number POMC and NPY positive cells in the ARH of offspring of obese mice was comparable to that of control mice (S1 Fig). During adult life, both the densities of POMC- and AgRP-labeled fibers were reduced (Fig 4B). Similar decreases in POMC and AgRP fiber densities were also observed in the adult DMH, which is another terminal field of ARH projections (S2 Fig).

**Fig 4.**
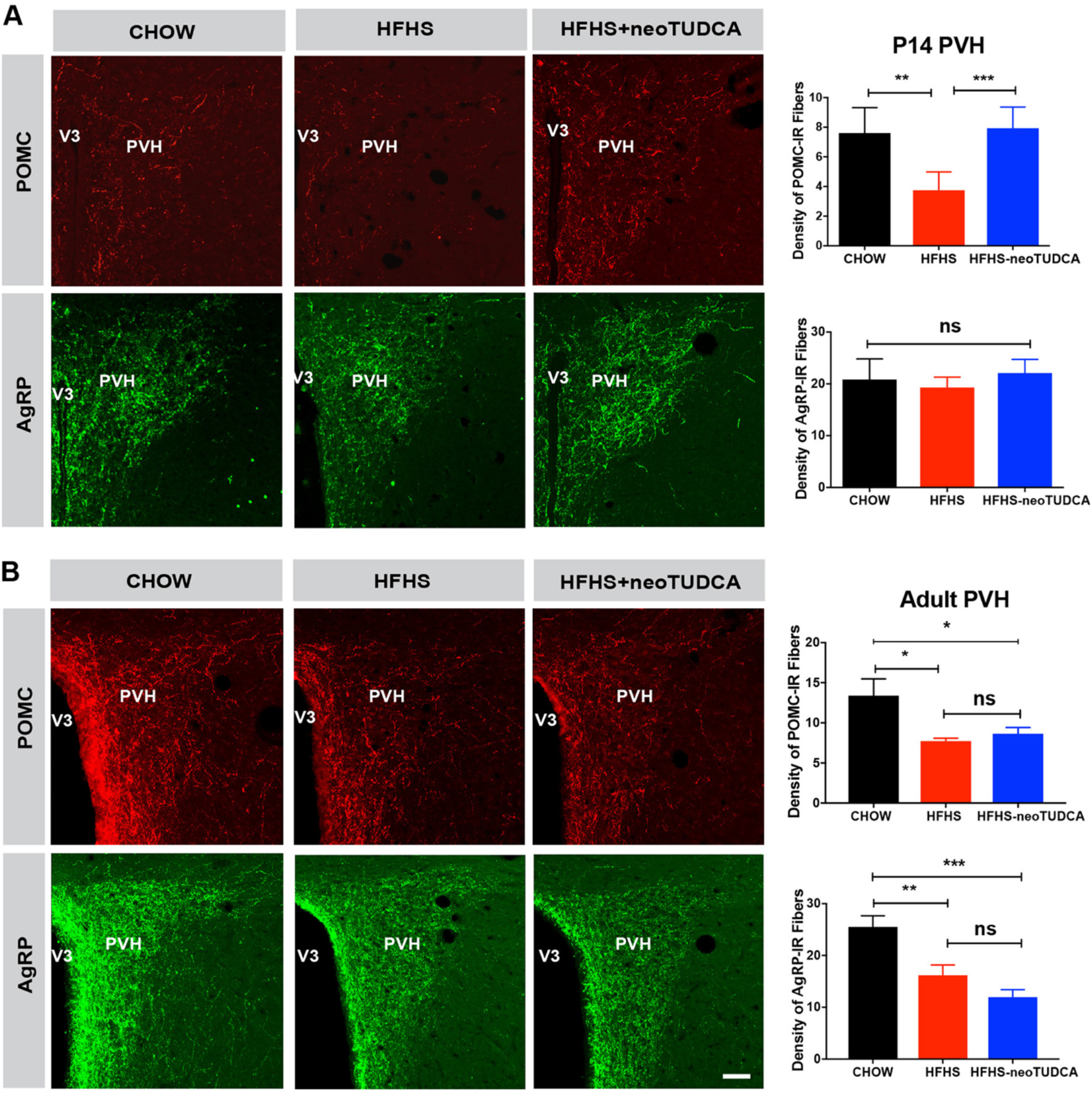
TUDCA treatment reverses neonatal disruption of POMC axonal projections induced by maternal obesity. Confocal images and quantification of the density of pro-opiomelanocortin (POMC)- and agouti-related peptide (AgRP)-immunoreactive fibers in the paraventricular nucleus (PVH) of (A) P14 and (B) 10- to 12-week-old mice born to chow-fed dams, HFHS-fed dams, or HFHS-fed dams and treated with TUDCA neonatally (n = 5-7 per group). Data are presented as mean ± SEM. \**P* ≤ 0.05, \*\**P* < 0.01, \*\*\**P* ≤ 0.001, and \*\*\*\**P* ≤ 0.0001 *versus* other groups. Statistical significance was determined by one-way ANOVA (A-B) followed by Tukey’s Multiple Comparison test. ARH, arcuate nucleus of the hypothalamus; PVH, paraventricular nucleus of the hypothalamus. Scale bar, 50 µm.

Because TUDCA treatment restores normal leptin signaling in the developing ARH, we also examined whether neonatal TUDCA treatment improved ARH projections. Neonatal injections of TUDCA in pups born to obese dams restored a normal density of POMC-labeled fibers in P14 pups born to obese dams (Fig 4A). However, enhancing ER capacity neonatally did not influence POMC or AgRP projections in adult animals (Fig 4B and S2 Fig)

### Maternal obesity increases circulating fatty acids concentration and treatment with saturated fatty acids induces ER stress and blunts ARH axonal outgrowth

Our results show that overconsumption of a western-style diet rich in fatty acids during pregnancy and lactation is associated with abnormal hypothalamic development. We also measured circulating fatty acid concentration during pregnancy and found that dams fed a HFHS diet have a 4-fold increase in serum fatty acid levels compared to control dams (Fig 5A). Offspring born to obese dams also displayed higher levels of circulating fatty acids at P10 that persisted into adulthood and neonatal TUDCA treatment restored normal levels of fatty acids (Fig 5A). To determine which type of fatty acids could cause neurodevelopmental abnormalities in our model, we reviewed the dietary fat content of the HFHS diet used in this study and found high concentrations (93.3%) of saturated fatty acids, including palmitic, lauric, and myristic acids and low concentrations (2.4%) of monounsaturated fats such as oleic acid (Table 1).

**Table 1.** Fatty acid composition of D12331 diet

| **Fatty acid** | **gm/kg diet** |
| --- | --- |
| C6, Caproic | 2.0 |
| C8, Caprylic | 25.7 |
| C10, Capric | 19.7 |
| C12, Lauric | 158.7 |
| C14, Myristic | 60.0 |
| C16, Palmitic | 31.6 |
| C18, Stearic | 36.3 |
| C18:1, Oleic | 8.7 |
| C18:2, Linoleic | 13.5 |
| C18:3, Linolenic | 2.0 |

**Fig 5.**
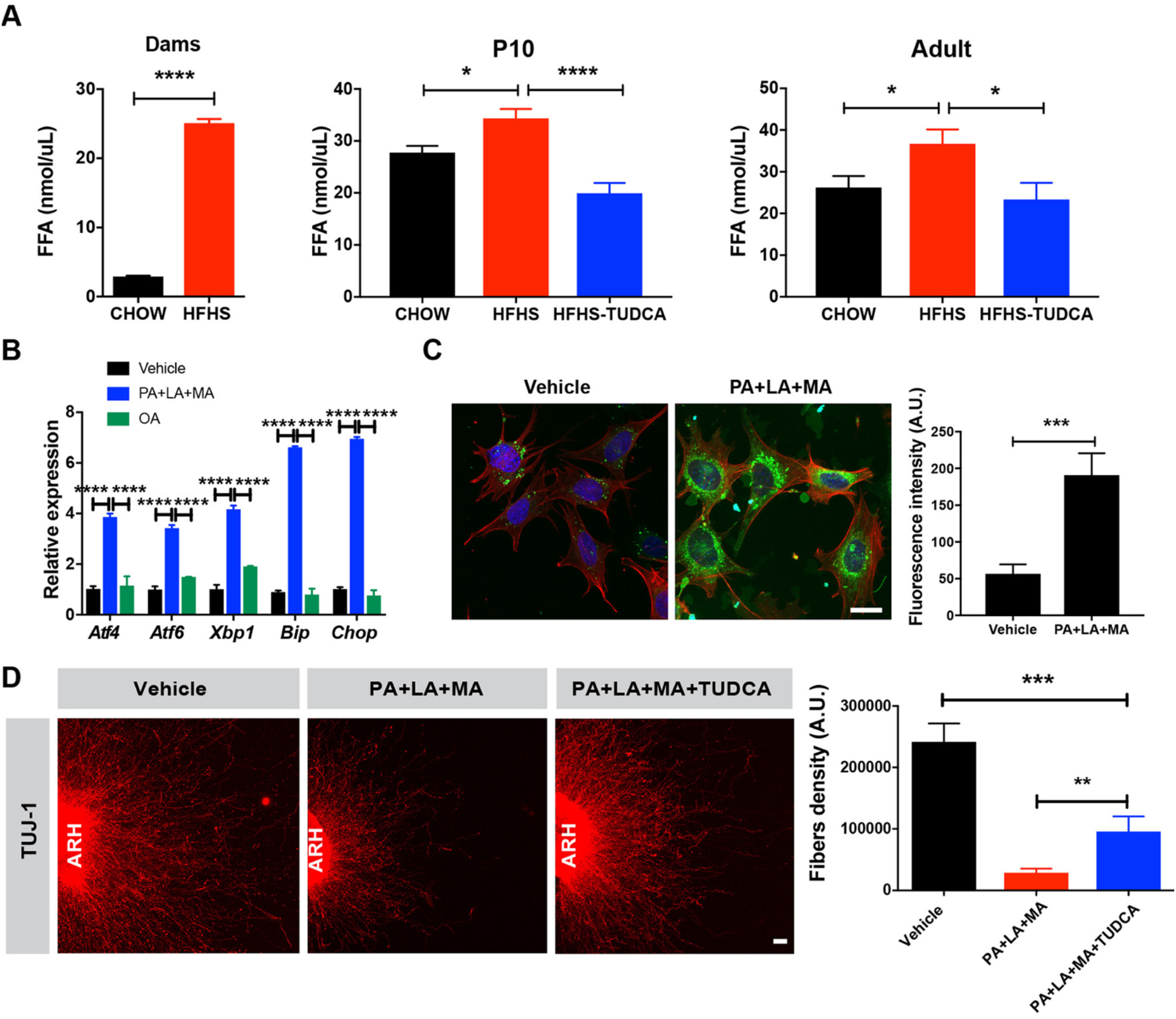
Saturated fatty acid treatment causes ER stress-induced disruption of axon growth. (A) Serum fatty acid levels in dams, P10 and 10-week-old mice born to chow-fed dams, HFHS-fed dams, or HFHS-fed dams and treated with TUDCA neonatally (n = 4-7 per group). (B) Relative expression of *Atf4, Atf6, Xbp1, Bip*, and *Chop* mRNA in mouse hypothalamic N43/5 cells treated with vehicle (BSA with 0.1% ethanol), or a cocktail of palmitate (250 µM) with lauric (1mM) and myristic acids (200 µM) (PA+LA+MA), or oleic acid alone (OA) for 24h (n = 4-5 cultures per condition). (C) Representative images and quantification of the density of long-chain fatty acid analog BODIPY (*green* fluorescence) immunoreactivity in N43/5 cells treated with vehicle (BSA with 0.1% ethanol) or palmitate (250 µM) with lauric (1mM) and myristic acids (200 µM) (PA+LA+MA) for 24h (n = 5-7 cultures per condition). Red fluorescence and blue fluorescence depict actin filaments phalloidin and DAPI nuclear staining, respectively. (D) Confocal images and quantification of TUJ1 (neuron-specific class III beta-tubulin) immunoreactive fibers derived from isolated arcuate nucleus (ARH) explants incubated with vehicle (0.1% ethanol) or a combination of palmitate (250 µM) with lauric (1mM) and myristic acids (200 µM) (PA+LA+MA) with or without TUDCA (750 µg/ml, n = 6 cultures per condition). Data are presented as mean ± SEM. \**P* < 0.05, \*\**P* ≤ 0.01, \*\*\**P* < 0.001 *versus* other groups. Statistical significance was determined by unpaired two-tailed Student’s t test (A, C, D), two-way ANOVA followed by Tukey’s Multiple Comparison test (B). Scale bars, 20 µm (C), and 50 µm (D).

Direct exposure of N43/5 cells to individual saturated fatty acids such as palmitic, lauric, or myristic acids or a combination of these fatty acids increased ER stress markers gene expression (S3 Fig). In particular, the mRNA expression of *Atf4, Atf6, Xbp1, Bip*, and *Chop* was 4- to 7-fold increased in cells treated with a combination of palmitic, lauric, and myristic acids compared to vehicle-treated cells (Fig 5B). In contrast, expression of ER stress markers was not affected when cells were treated with the monousaturated fat oleic acid (Fig 5B).

We next assessed fatty acids intracellular transport in hypothalamic cells using BODIPY, a fluorescent long-chain fatty acids analog. Exposure of hypothalamic N43/5 cells to a combination of palmitic, lauric, and myristic acids resulted in greater BODIPY labeling in N43/5 cell bodies compared to vehicle-treated cells (Fig 5C). In order to determine if these saturated fatty acids also impacted ARH axon growth and whether it involves ER stress, we also performed a series of *in vitro* experiments in which ARH explants were microdissected, placed in a collagen matrix, and then exposed to combination of saturated fatty acids (*i.e.*, palmitic, lauric, and myristic acids), or saturated fatty acids with TUDCA, or vehicle alone. After 48 hours, the density of TUJ1-labeled neurite, neuron-specific class III beta-tubulin, from ARH explants treated with saturated fatty acids was approximately 10-fold lower than that of vehicle-treated explants (Fig 5D). Moreover, pre-incubation of ARH explants with TUDCA improved disrupted axon outgrowth after saturated fatty acids treatment (Fig 5D).

Together, these data indicate that maternal obesity caused elevated circulating fatty acid levels in the dams and offspring and that direct exposure to saturated fatty acids induced ER stress gene expression in hypothalamic cells. They also show that saturated fatty acids can be transported in hypothalamic cells blunting axon growth, and that this phenomenon appears to involve ER stress pathways.

## Discussion

Although the link between perinatal overnutrition and lifelong metabolic regulation has been clearly shown, little is known about the mechanisms underlying this programming effect. In this study, we show that maternal obesity causes lifelong metabolic alterations associated with abnormal development of hypothalamic feeding circuits in the offspring. We also report that maternal obesity induces ER stress in key tissues involved in energy metabolism during critical periods of growth and development, particularly in the arcuate nucleus and pancreas. Moreover, we found that relieving ER stress neonatally ameliorates metabolic and hypothalamic structural abnormalities in animals born to obese dams and that these effects are likely mediated through increased leptin sensitivity. Furthermore, we report that the malprogramming action of maternal obesity on hypothalamic development involves a direct effect of saturated fatty acids on arcuate axon growth.

Our findings are generally consistent with previous works showing that maternal obesity causes lifelong weight gain and glucose intolerance associated with disruption in AgRP/NPY and POMC axonal projections during adult life [10,11]. However, our study reveals that maternal obesity does not affect, during early postnatal life, AgRP circuits whereas it affects POMC axonal projections, suggesting that distinct mechanisms underlie the effects of maternal HFHS feeding on POMC *versus* AgRP/NPY neurons. Vogt and colleagues have specifically attempted to compare the consequences of maternal obesity during gestation and lactation and have shown that maternal consumption of HFD during lactation (but not during pregnancy) is sufficient to cause obesity and diabetes and to alter the development of POMC projections in the offspring [10]. Consistent with the importance of the postnatal period in the nutritional programming of metabolism and hypothalamic circuits, exposure to chronic postnatal overnutrition by rearing neonates in small litters also predisposes to obesity and disrupts hypothalamic development [22,23,24].

A variety of developmental pathways control the development of arcuate feeding circuits. Among this array of signals, attention has been given to leptin. The density of ARH axonal projections is reduced in leptin-deficient mouse neonates and adults, which can be rescued with leptin treatment during early postnatal life [8,25,26]. Moreover, leptin appears to exert its neurodevelopmental actions on arcuate circuits though LepRb→pSTAT3 signaling [21]. The data presented here indicate that maternal obesity causes chronic hyperleptinemia in the offspring associated with reduced arcuate leptin-induced pSTAT3 during a critical period of hypothalamic development. A similar increase in circulating leptin levels and a reduction in arcuate leptin sensitivity has been reported in rat neonates exposed to chronic postnatal overnutrition [22]. However, the mechanisms underlying this early leptin resistance remained elusive. Here, we show that relieving ER stress enhances arcuate leptin resistance and improves hypothalamic development and long-term metabolic outcomes. These findings are consistent with previous data showing that ER stress inducers, such as tunicamycin, blunts neurite elongation and induce a collapse of neuronal growth cones from PC-12 cells or dissociated rat sensory neurons [27]. The site of action of TUCDA remains to be determined but it likely involves a direct effect on ARH neurons. The highest level of ER stress induction is observed in the ARH and neonatal TUDCA treatment normalizes arcuate ER stress gene expression. Moreover, previous studies have reported that the pharmacological induction and genetic loss of the ER stress function in the brain block hypothalamic leptin-induced STAT3 activation [14].

Future studies are needed to determine the contribution of specific ER stress pathways to the nutritional programming of hypothalamic development and leptin resistance. However, previous studies have shown that overexpression of *Xbp1* or *Atf6* in mouse embryonic fibroblast cells increases their resistance to the inhibitory effects of tunicamycin and prevents ER stress-mediated inhibition of leptin signaling [14]. Moreover, when fed a high fat diet, mice lacking *Xbp1* in neurons display an obesogenic phenotype, associated with hyperphagia and reduced oxygen consumption [14]. In addition, leptin-induced STAT3 phosphorylation is significantly attenuated in the hypothalamus of these mice [14]. Furthermore, constitutive expression of a dominant *Xbp1* form specifically in POMC neurons leads to a lean phenotype, characterized by increased energy expenditure and leptin sensitivity, further supporting a fundamental role for the XBP1 pathway in POMC neurons in the deleterious metabolic effects of hypothalamic ER stress [15].

Our results show that lipid overload, especially saturated fatty acids, triggers ER stress in hypothalamic cells and that it contributes to disruption in arcuate axon growth. Previous studies have demonstrated that hypothalamic neurons can sense circulating fatty acids, and that endogenous lipid metabolism in the hypothalamus is a key mechanism regulating whole-body energy balance [28]. Adult rodents fed a HFD exhibit elevated concentrations of fatty acids in the hypothalamus, which causes an accumulation of palmitoyl-CoA and other harmful species [29,30]. Moreover, studies in hypothalamic cell lines have demonstrated that palmitate triggers ER stress and apoptosis [31,32,33]. In addition, intracerebroventricular injection of saturated fatty acids *in vivo* induces ER stress in the hypothalamus of rats [34]. Notably, palmitate decreases protein abundance and function of the α-MSH receptor MC4-R and chemical chaperone reverses this biochemical abnormality [35], suggesting that saturated fatty acids may not only cause disruption in the development of POMC axonal projections, but also attenuate the post-synaptic action of POMC-derived peptides through a ER stress-dependent mechanism.

## Methods

### Animals

All animal procedures were conducted in compliance with and approved by the IACUC of the Saban Research Institute of the Children’s Hospital of Los Angeles. The animals were housed under specific pathogen-free conditions, maintained in a temperature-controlled room with a 12 h light/dark cycle, and provided *ad libitum* access to water and standard laboratory chow (Special Diet Services). At 7 weeks of age, female C57BL/6J wild-type (WT) mice were placed on either a regular chow diet [4.5 kcal% fat, provided by PicoLab Rodent Diet 5053] or a high-fat, high-sugar (HFHS) diet [58 kcal% fat with sucrose, provided by Research Diet D12331] for 6 weeks before mating. The mice were kept on their respective diets throughout pregnancy and lactation. Male breeders were fed a normal chow diet. Offspring were fed a normal chow diet after weaning. Litter sizes were standardized to six pups 48 hours post-delivery, and attempts were made to maintain an equal sex ratio. Only male mice were studied.

### Neonatal TUDCA treatment

The mice were injected intraperitoneally daily with TUDCA (Millipore, 150 mg/kg/day) from P4 to P16. Controls received injections with an equivalent volume of vehicle (0.9% NaCl).

### Tissue collection

The ARH and PVH of P10 and 10-week-old mice were dissected under a stereomicroscope. Liver, pancreas and epididymal white adipose tissues were collected from P10 and 10-week-old mice.

### Cell culture and fatty acid treatment

The embryonic mouse hypothalamic cell line N43/5 was cultured in Dulbecco’s modified Eagle’s medium (Sigma, D5796) supplemented with 10% fetal bovine serum, 100 U/ml penicillin and 100 µg/ml streptomycin at 37°C in 5% CO_2_ and a humidified atmosphere. N43/5 cells were plated out at density of 6×10^5^ cells per well in a 6-wells plate. The following day, medium was changed to culture medium containing either vehicle (BSA with 0.1% ethanol; Sigma), or palmitic (PA; 250 µM; Sigma), lauric (LA; 1mM; Sigma), myristic (MA; 200 µM; Sigma), or oleic acids (OA; 250 µM; Sigma), or combination of these fatty acids for 24h.

### RNA extraction and RT-qPCR analyses

Total RNA was isolated using the Arcturus PicoPure RNA Isolation Kit (for hypothalamic samples) (Life Technologies), the RNeasy Lipid Tissue Kit (for peripheral samples) (Qiagen), or PureLink RNA mini kit (for N43/5 cell samples). cDNA was generated with the High-Capacity cDNA Reverse Transcription Kit (Life Technologies). Quantitative real-time PCR was performed using TaqMan Fast Universal PCR Master Mix and the commercially available TaqMan gene expression primers: *Atf4* (Mm00515324_m1), *Atf6* (Mm01295317_m1), *Xbp1* (Mm00457357_m1) *Bip* (Mm00517691_m1), *Chop* (Mm00492097_m1), and *Gapdh* (Mm99999915_g1). mRNA expression was calculated using the 2^-ΔΔCt^ method after normalization to the expression of the *Gapdh* housekeeping gene. All assays were performed using an Applied Biosystems 7900 HT real-time PCR system.

### Physiological measures

The maternal body weight was recorded weekly until the end of pregnancy. Offspring (n = 5 per group) was weighed weekly 1 to 10 weeks of age using an analytical balance. Body composition analysis (fat/lean mass) was performed in pregnant females at gestational day 16 and in the offspring at 10 weeks of age using NMR (Echo MRI). Food intake, O_2_ and CO_2_ production, energy expenditure, respiratory exchange ratio (*i.e.*, VCO_2_/O_2_), and locomotor activity (XY) were monitored at 10 weeks of age using a combined indirect calorimetry system (TSE Systems). The mice were acclimated in monitoring chambers for 2 days, and the data were collected for 3 days. These physiological measures were performed at the Rodent Metabolic Core of Children’s Hospital of Los Angeles.

Glucose and insulin tolerance tests (GTT and ITT) were conducted in 7-8-week-old mice through i.p. injection of glucose (1.5 mg/g body weight) or insulin (2U/kg body weight) after overnight fasting. Blood glucose levels were measured at 0, 15, 30, 45, 60, 90, 120, and 150 min post-injection, as previously described [36].

Serum leptin levels were assayed in chow-fed or HFHS-fed mothers at gestational day 16, and in the offspring of chow- or HFHS-fed dams at E16.5, P10 and 10 weeks of age using a commercially available leptin ELISA kit (Millipore). Serum free fatty acid levels were assayed in chow-fed or HFHS-fed mothers at gestational day 16 and in the offspring of chow- or HFHS-fed dams at P10 and 10 weeks of age using a commercially available FFA kit (Abcam).

### POMC, AgRP, and NPY immunohistochemistry

Ten- to 12-week-old mice were perfused transcardially with 4% paraformaldehyde. The brains were then frozen, sectioned at 30-µm thick, and processed for immunofluorescence using standard procedures [8,37]. The primary antibodies used for IHC were as follows: rabbit anti-POMC (1:20,000, Phoenix Pharmaceuticals), rabbit anti-AgRP (1:1,000, Phoenix Pharmaceuticals), and sheep anti-NPY (1:3,000, Abcam). The primary antibodies were visualized with Alexa Fluor 647 donkey anti-sheep IgG, Alexa Fluor 488 donkey anti-rabbit IgG, or Alexa Fluor 488 donkey anti-mouse IgG, or Alexa Fluor 568 donkey anti-rabbit IgG (1:200, Millipore). The sections were counterstained using bis-benzamide (1:10,000, Invitrogen) to visualize cell nuclei.

### pSTAT3 immunohistochemistry

Leptin (3 mg/kg; Peprotech) was injected intraperitoneally in P14 pups. Animals were perfused 45 min later with a solution of 2% paraformaldehyde. Frozen coronal sections were cut at 30 µm and pretreated for 20 min in 0.5% NaOH and 0.5%H_2_O_2_ in KPBS, followed by immersion in 0.3% glycine for 10 min. Sections were then placed in 0.03% SDS for 10 min and placed in 4% normal serum + 0.4% Triton X-100 + 1% BSA (fraction V) for 20 min before incubation for 48h with a rabbit anti-pSTAT3 antibody (1:1,000, Cell Signaling). The primary antibody was localized with Alexa Fluor 568 Goat anti-Rabbit IgGs (Invitrogen; 1:200). Sections were counterstained using bis-benzamide (Invitrogen; 1:10,000) to visualize cell nuclei, and coverslipped with buffered glycerol (pH 8.5).

### Histomorphological assessment of white adipose tissue

Epididymal white adipose tissue from 10-weekold mice was collected, fixed in a 4% paraformaldehyde solution, sectioned at 5 µm, and then stained with a Perilipin A antibody (1:1,000, Sigma) using standard procedures.

### BODIPY staining

N43/5 cells were treated with vehicle or a combination of palmitate (250 µM; Sigma) with lauric (1mM; Sigma) and myristic acids (200 µM; Sigma) for 24 hr and 2 µM of Bodipy 493/503 (4,4-difluro-1,3,5,7-tetramethyl-4-bora-3a,4a-diaza-s-indacene; Invitrogen) and Alexa Fluor 568 Phalloidin (0.1 µM; Invitrogen) were added to culture media for 15 min at room temperature. N43/5 were then fixed in a solution of 4% paraformaldehyde for 5 min and washed with KPBS. Slides were counterstained using bis-benzamide (Invitrogen; 1:10,000) to visualize cell nuclei.

### Isolated ARH explant culture

Brains were collected from P4 mice and sectioned at a 200-µm thickness with a vibroslicer as previously described [8,37]. The ARH was then carefully dissected out of each section under a stereomicroscope. Explants were cultured onto a rat tail collagen matrix (BD Bioscience) and each explant was pre-treated for 6 h with fresh modified Basal Medium Eagle (Invitrogen) containing TUDCA (750µg/ml) or vehicle followed by a combination of palmitate (250 µM) with lauric (1mM) and myristic acids (200 µM) or vehicle alone (BSA with 0.1% ethanol). After 48 h, the explants were fixed in paraformaldehyde and neurites extending from the explants were stained with TUJ1 (β III tubulin) (rabbit, 1:5,000, Covance) as described previously [37].

### Image analysis

The images were acquired using a Zeiss LSM 710 confocal system equipped with a 20X objective through the ARH (for cell numbers), through the PVH and the DMH (for fibers density), through adipose tissue (for adipocyte size), and through N43/5 cell cultures (for BODIPY staining). The average number of cells and density of fibers were analyzed in 2-4 sections per culture. For the explant experiments, sections were acquired using a Zeiss LSM 710 confocal system equipped with a 10X objective. Slides were numerically coded to obscure the treatment group. The image analysis was performed using ImageJ analysis software (NIH) as previously described [21,36,37].

For the quantitative analysis of cell number, POMC^+^, NPY^+^, and pSTAT3^+^ cells were manually counted. Only cells with corresponding bis-benzamide-stained nuclei were included in our counts.

Determination of mean adipocyte size (µm^2^) was measured semi-automatically using the FIJI distribution [30] of Image J software (NIH, ImageJ1.47i). The average adipocyte size measured from 3 fields and six sections in each mouse was used for statistical comparisons used for statistical comparisons.

For the quantitative analysis of fiber density (for POMC, AgRP, and TUJ1 fibers) and BODIPY fluorescence, each image plane was binarized to isolate labeled materials from the background and to compensate for differences in fluorescence intensity. The integrated intensity, which reflects the total number of pixels in the binarized image, was then calculated for each image as previously described [8,37]. This procedure was conducted for each image plane in the stack, and the values for all of the image planes in a stack were summed.

### Statistical analysis

All values are represented as the mean ± SEM. Statistical analyses were conducted using GraphPad Prism (version 5.0a). Data sets with only two independent groups were analyzed for statistical significance using unpaired two-tailed Student’s t test. Data sets with more than two groups were analyzed using one-way analysis of variance (ANOVA) followed by the Tukey’s Multiple Comparison test. For statistical analyses of body weight, GTT ITT, and RER, we performed two-way ANOVAs followed by Tukey’s Multiple Comparison test. Statistically significant outliers were calculated using Grubb’s test for outliers. P ≤ 0.05 was considered statistically significant.

## Supplementary material

Supplementary material includes 3 figures and can be found with this article online.

## Acknowledgments

We thank Brad Wanken and the CHLA Rodent Metabolic Core for metabolic studies. We also thank the CHLA Cellular Imaging Core for confocal imaging studies. We are also grateful to Gricelda Vasquez for her assistance with animal husbandry. This study was supported by the National Institutes of Health (Grants DK84142, DK102780, and DK118401 to SGB).

## Author Contributions

S.P. conceived, designed, and performed most of the experiments and analyzed the data. A.J. performed and analyzed some of the immunohistochemical experiments. S.G.B. conceived, designed, and supervised the project. S.P. and S.G.B. wrote the manuscript.

## References

1. McMillen IC, Adam CL, Muhlhausler BS (2005) Early origins of obesity: programming the appetite regulatory system. J Physiol (Lond) 565: 9-17.

2. Bouret S, Levin BE, Ozanne SE (2015) Gene-Environment Interactions Controlling Energy and Glucose Homeostasis and the Developmental Origins of Obesity. 47-82 p.

3. Sullivan EL, Grove KL (2010) Metabolic imprinting in obesity. Forum Nutr 63: 186-194.

4. Taylor PD, Poston L (2007) Developmental programming of obesity in mammals. Exp Physiol 92: 287-298.

5. Martin-Gronert MS, Ozanne SE (2005) Programming of appetite and type 2 diabetes. Early Human Development 81: 981-988.

6. Horvath TL, Bruning JC (2006) Developmental programming of the hypothalamus: a matter of fat. Nat Med 12: 52-53.

7. Chen H, Simar D, Morris MJ (2009) Hypothalamic Neuroendocrine Circuitry is Programmed by Maternal Obesity: Interaction with Postnatal Nutritional Environment. PLoS ONE 4: e6259.

8. Bouret SG, Draper SJ, Simerly RB (2004) Trophic Action of Leptin on Hypothalamic Neurons That Regulate Feeding. Science 304: 108-110.

9. Bouret SG, Gorski JN, Patterson CM, Chen S, Levin BE, et al. (2008) Hypothalamic Neural Projections Are Permanently Disrupted in Diet-Induced Obese Rats. Cell Metabolism 7: 179-185.

10. Vogt MC, Paeger L, Hess S, Steculorum SM, Awazawa M, et al. (2014) Neonatal Insulin Action Impairs Hypothalamic Neurocircuit Formation in Response to Maternal High-Fat Feeding. Cell 156: 495-509.

11. Kirk SL, Samuelsson A-M, Argenton M, Dhonye H, Kalamatianos T, et al. (2009) Maternal Obesity Induced by Diet in Rats Permanently Influences Central Processes Regulating Food Intake in Offspring. PLoS ONE 4: e5870.

12. Ozcan U, Cao Q, Yilmaz E, Lee A-H, Iwakoshi NN, et al. (2004) Endoplasmic Reticulum Stress Links Obesity, Insulin Action, and Type 2 Diabetes. Science 306: 457-461.

13. Özcan U, Yilmaz E, Özcan L, Furuhashi M, Vaillancourt E, et al. (2006) Chemical Chaperones Reduce ER Stress and Restore Glucose Homeostasis in a Mouse Model of Type 2 Diabetes. Science 313: 1137-1140.

14. Ozcan L, Ergin AS, Lu A, Chung J, Sarkar S, et al. (2009) Endoplasmic reticulum stress plays a central role in development of leptin resistance. Cell Metab 9: 35-51.

15. Williams KW, Liu T, Kong X, Fukuda M, Deng Y, et al. (2014) Xbp1s in Pomc neurons connects ER stress with energy balance and glucose homeostasis. Cell Metab 20: 471-482.

16. Perlmutter DH (2002) Chemical chaperones: a pharmacological strategy for disorders of protein folding and trafficking. Pediatr Res 52: 832-836.

17. Attig L, Solomon G, Ferezou J, Abdennebi-Najar L, Taouis M, et al. (2008) Early postnatal leptin blockage leads to a long-term leptin resistance and susceptibility to diet-induced obesity in rats. Int J Obes 32: 1153-1160.

18. Yura S, Itoh H, Sagawa N, Yamamoto H, Masuzaki H, et al. (2005) Role of premature leptin surge in obesity resulting from intrauterine undernutrition. Cell Metabolism 1: 371-378.

19. Vickers MH, Gluckman PD, Coveny AH, Hofman PL, Cutfield WS, et al. (2005) Neonatal Leptin Treatment Reverses Developmental Programming. Endocrinology 146: 4211-4216.

20. Vickers MH, Gluckman PD, Coveny AH, Hofman PL, Cutfield WS, et al. (2008) The Effect of Neonatal Leptin Treatment on Postnatal Weight Gain in Male Rats Is Dependent on Maternal Nutritional Status during Pregnancy. Endocrinology 149: 1906-1913.

21. Bouret SG, Bates SH, Chen S, Myers MG, Simerly RB (2012) Distinct Roles for Specific Leptin Receptor Signals in the Development of Hypothalamic Feeding Circuits. The Journal of Neuroscience 32: 1244-1252.

22. Glavas MM, Kirigiti MA, Xiao XQ, Enriori PJ, Fisher SK, et al. (2010) Early Overnutrition Results in Early-Onset Arcuate Leptin Resistance and Increased Sensitivity to High-Fat Diet. Endocrinology 151: 1598-1610.

23. Collden G, Balland E, Parkash J, Caron E, Langlet F, et al. (2015) Neonatal overnutrition causes early alterations in the central response to peripheral ghrelin. Molecular Metabolism 4: 15-24.

24. Caron E, Ciofi P, Prevot V, Bouret SG (2012) Alteration in Neonatal Nutrition Causes Perturbations in Hypothalamic Neural Circuits Controlling Reproductive Function. J Neurosci in press.

25. Bouyer K, Simerly RB (2013) Neonatal Leptin Exposure Specifies Innervation of Presympathetic Hypothalamic Neurons and Improves the Metabolic Status of Leptin-Deficient Mice. The Journal of Neuroscience 33: 840-851.

26. Kamitakahara A, Bouyer K, Wang CH, Simerly R (2017) A critical period for the trophic actions of leptin on AgRP neurons in the arcuate nucleus of the hypothalamus. J Comp Neurol.

27. Patterson S, Skene J (1994) Novel inhibitory action of tunicamycin homologues suggests a role for dynamic protein fatty acylation in growth cone-mediated neurite extension. The Journal of Cell Biology 124: 521-536.

28. Martinez de Morentin PB, Varela L, Ferno J, Nogueiras R, Dieguez C, et al. (2010) Hypothalamic lipotoxicity and the metabolic syndrome. Biochim Biophys Acta 1801: 350-361.

29. Benoit SC, Kemp CJ, Elias CF, Abplanalp W, Herman JP, et al. (2009) Palmitic acid mediates hypothalamic insulin resistance by altering PKC-theta subcellular localization in rodents. J Clin Invest 119: 2577-2589.

30. Posey KA, Clegg DJ, Printz RL, Byun J, Morton GJ, et al. (2009) Hypothalamic proinflammatory lipid accumulation, inflammation, and insulin resistance in rats fed a high-fat diet. Am J Physiol Endocrinol Metab 296: E1003-1012.

31. Mayer CM, Belsham DD (2010) Palmitate attenuates insulin signaling and induces endoplasmic reticulum stress and apoptosis in hypothalamic neurons: rescue of resistance and apoptosis through adenosine 5’ monophosphate-activated protein kinase activation. Endocrinology 151: 576-585.

32. Choi SJ, Kim F, Schwartz MW, Wisse BE (2010) Cultured hypothalamic neurons are resistant to inflammation and insulin resistance induced by saturated fatty acids. Am J Physiol Endocrinol Metab 298: E1122-1130.

33. McFadden JW, Aja S, Li Q, Bandaru VV, Kim EK, et al. (2014) Increasing fatty acid oxidation remodels the hypothalamic neurometabolome to mitigate stress and inflammation. PLoS One 9: e115642.

34. Milanski M, Degasperi G, Coope A, Morari J, Denis R, et al. (2009) Saturated fatty acids produce an inflammatory response predominantly through the activation of TLR4 signaling in hypothalamus: implications for the pathogenesis of obesity. J Neurosci 29: 359-370.

35. Cragle FK, Baldini G (2014) Mild lipid stress induces profound loss of MC4R protein abundance and function. Mol Endocrinol 28: 357-367.

36. Coupe B, Ishii Y, Dietrich MO, Komatsu M, Horvath TL, et al. (2012) Loss of Autophagy in Pro-opiomelanocortin Neurons Perturbs Axon Growth and Causes Metabolic Dysregulation. Cell Metabolism 15: 247-255.

37. Steculorum SM CG, Coupe B, Croizier S, Andrews Z, Jarosch F, Klussmann S, Bouret SG. (2015) Ghrelin programs development of hypothalamic feeding circuits.. The Journal of Clinical Investigation 125: 846–858.

